# Dynamic Sex Differences in Appetitive and Reactive Aggression

**DOI:** 10.1101/2022.02.22.481480

**Authors:** Antonio Aubry, C. Joseph Burnett, Nastacia L. Goodwin, Long Li, Jovana Navarrete, Yizhe Zhang, Valerie Tsai, Romain Durand-de Cuttoli, Sam A. Golden, Scott J. Russo

## Abstract

Aggression is an evolutionarily conserved, adaptive component of social behavior. Studies in male mice illustrate that aggression is influenced by numerous factors including the degree to which an individual finds aggression rewarding and will work for access to attack and subordinate mice. While such studies have expanded our understanding of the molecular and circuit mechanisms of male aggression very little is known about female aggression, owed in part to limited availability of valid mouse models in females. Here we use an ethologically relevant model of male vs. female aggression by pair housing adult male and female outbred CFW mice with opposite sex cage mates. We assess reactive (defensive) aggression in the resident intruder (RI) test and appetitive (rewarding) aggression in the aggression conditioned place preference (CPP) and operant self-administration (SA) tests. Our results show dramatic sex differences in both qualitative and quantitative aspects of reactive vs. appetitive aggression. Males exhibit more wrestling and less investigative behavior during RI, find aggression rewarding and will work for access to a subordinate to attack. Females exhibit more bites, alternate between aggressive behaviors and investigative behaviors more readily during RI, however, they do not find aggression to be rewarding or reinforcing. These results establish sex differences in aggression in mice, providing an important resource for the field to better understand the circuit and molecular mechanisms of aggression in both sexes.

## Introduction

Aggression exists along a spectrum from adaptive to maladaptive and is governed by both reactive (defensive) and appetitive (rewarding) drives. The transition away from an adaptive state can be associated with neuropsychiatric conditions and presents a challenge to patients and caregivers. Modeling and understanding the behavioral etiology of aggressive behavior is therefore a health priority with the potential to guide therapeutic interventions across a number of neuropsychiatric diseases. In mice, aggressive behavior serves as an evolutionary adaptation for survival [1] and engages highly conserved neural mechanisms [2]. However, while aggression is often a focus of both popular and scientific inquiry and highly evolutionarily conserved, very little is known about the neural and behavioral mechanisms controlling aggression-related sex differences.

Recently, preclinical behavioral models have been introduced that facilitate the direct comparison of reactive and appetitive aggression [3–5]. Typically, reactive aggression is investigated using the resident intruder (RI) test in which a male intruder is introduced to the home cage of a male resident and they are allowed to freely interact [6]. RI testing addresses territorial and/or reactive aggression but is unable to dissociate reactive vs. appetitive components. To assess aggression reward, aggression conditioned place preference (CPP) can be used where male mice will display a preference for contexts previously associated with opportunities to attack a naïve conspecific [7–9]. However, like RI testing, this procedure uses forced involuntary social interactions and therefore cannot fully dissociate reactive from appetitive components. To overcome this, several groups have developed social operant tasks that measure voluntary appetitive aggression seeking in male mice [10–13], and established that appetitive aggression can transition to compulsive addiction-like behavior in some mice (Golden et al., 2017). These behavioral advances provide a toolbox for the exploration of shared and dissociable neural mechanisms underlying the spectrum of aggression behaviors.

However, while these procedures have proven effective for studying aggressive behavior and the underlying neurobiology in animal models, studies have focused nearly entirely on males. Male aggression is typically assessed in the context of isolated housing, and under similar conditions naïve female mice do not show comparable intruder-directed aggression. To overcome this, alternative model organisms can be used such as Syrian hamsters (*Mesocricetus auratus*)[14] and California mice (*Peromyscus californicus*)[15], or with mouse models of gestational aggression [16–18]. However, there are limitations to these alternate models: (*i*) only *Mus musculus* models can presently exploit the broad transgenic toolbox available to understand circuit and molecular mechanisms and (*ii*) gestational aggression models are not ideal for evaluating sex differences in aggressive behavior since these behaviors are linked to hormonal changes specifically associated with pregnancy, parturition and lactation.

Two recent studies have revisited female aggression during RI tests using outbred mouse strains, as opposed to the more typically used inbred strains, and have found that naïve outbred CFW mice can display similar levels of aggression as males. Outbred mouse strains such as CD1 and CFW have gained popularity in aggression-related studies due to their high individual variability in innate aggressive behavior (Chia et al., 2005, Golden et al., 2016). Similarly, isolated sex-naive female outbred CFW, but not inbred female C57BL6/J, mice will attack juvenile male or adult female C57BL6/J intruders in the home cage [19], and female CFW mice pair-housed with a castrated male partner will attack adult female C57BL6/J mice, with similar but non-identical behavioral strategies to males (Newman et al. 2019). The establishment of a female RI procedure opens the door for sex comparisons of the neurobiological substrates of reactive aggression, but currently there are no direct comparisons of appetitive aggression in males vs. females.

To further extend preclinical models of aggression, we directly compared adult male and female CFW mice in reactive and appetitive aggression procedures to evaluate sex as a biological variable. The evaluation of sex differences in complex social behaviors, such as aggression, are compounded by the need for detailed ethological annotation of behavioral actions and sequences. Typically, only simple statistics of aggression, including number of bouts, latency to first attack, and total attack duration, are reported. This lack of in-depth behavioral analysis obscures potential differences in specific attack behaviors and behavioral sequences and prevents an understanding of the role of reciprocal social interactions in driving or preventing attacks.

A recent study [20] utilized confirmatory factor analysis to identify individual differences in aggression, using a small selection of typically reported behaviors – aggression latency, bouts, durations – to generate easily generalizable behavioral models for improved reporting of aggression variability in social defeat behavior across research sites. Building on this effort, and to extend such analysis for more granular behavioral features, we utilized a discrete state hidden Markov model (HMM). HMMs analyze the ordering, clustering, and transitions between actions and are able to cluster time-series of behavior into distinct hidden states. Each state is associated with an emission probability matrix which represents how likely a given behavior is to occur while and animal is in a given state. These models have been used extensively in analyzing sequences of speech, gestures, animal behavior and the analysis of gene and protein sequences [21–24]. The discrete state HMM allowed us to examine the temporal composition of social behaviors and the hidden states which contribute to male versus female aggression.

We therefore set out in this study to accomplish two goals: (*i*) To examine whether or not males and females show different suites of behavior or different temporal arrangements of aggressive and investigative behavior throughout reactive aggression encounters and (*ii*) to compare reactive and appetitive aggression between male and female mice.

While we found no difference in simple readouts of aggression behavior in resident intruder assays (i.e., attack duration and latency to attack in aggressive mice), we observed clear sex differences in the behavioral sequences that make up bouts of aggressive and investigative behavior. The HMM revealed that females are more likely to switch between aggressive and investigative behaviors within a given interaction bout. In contrast, males tend to engage in interaction bouts that consist solely of one of the two types of social behavior (aggressive or investigative).

Further, using both aggression CPP and operant social self-administration (SA) procedures, we show that female mice, regardless of aggression expression during RI testing, do not exhibit context-dependent aggression reward nor do they exhibit appetitive aggression seeking behavior. To our knowledge, this is the first report of sex differences in appetitive vs. reactive aggression in *mus musculus*. Together these data support distinct patterns of aggressive behavior between males and female outbred mice and underscore the importance of future research to identify detailed mechanisms by which different sexes express aggressive behavior.

## Methods

### Mice

10-week-old CFW mice (Charles River Laboratories) were used as experimental mice for all studies. For studies conducted at Mount Sinai, females were pair-housed with a castrated male for the study duration (Newman et al., 2019) and males were pair-housed for 48-hr with stimulus females prior to isolate housing. Subject males were separated from group-housed cage mates and paired with a female for 2-d, then singly housed for an additional 10-d before all protocols. This procedure was used in order to acquire roughly equal amounts of aggressors (AGG) and non-aggressors (NON) without affecting the amount of aggression observed in AGGs. Subject female mice were housed with surgically castrated male CFW mice (see Castration procedure below) for at least 14-d before all protocols. 12-week-old C57BL/6J mice were used as intruders for all social interactions. All studies were conducted during the light cycle. Procedures were performed in accordance with the National Institutes of Health Guide for Care and Use of Laboratory Mice and approved by the Icahn School of Medicine at Mount Sinai Institutional Animal Care and Use Committee.

For studies conducted at the University of Washington, male and female CFW mice were pair-housed using an identical procedure as described above. 8-10 week-old C57BL/6J mice were used as intruders for all social interactions. Additional groups of two-week isolated males and females were also used as subject mice to assess the temporal effects of housing conditions. All studies were conducted during the dark cycle. Procedures were performed in accordance with the National Institutes of Health Guide for Care Use of Laboratory Mice and approved by the University of Washington Institutional Animal Care and Use Committee.

### Castration

Surgical castration was performed in-house as previously described [25]. Briefly, male mice were anesthetized by intraperitoneal injection with a mixture of ketamine HCl (100 mg/kg) and xylazine (10 mg/kg). Incisions were made in skin and peritoneum above the abdomen. The testes and testicular fat pads were extracted from the peritoneum. Testicular artery and fat pad were slowly cauterized using a high-temperature cautery pen (Bovie Medical Corporation). The peritoneum was then sealed with absorbable sutures and the skin stapled closed. Mice were group-housed for at least one week before pairing with female mice. At the University of Washington, surgical castration was conducted under inhaled isoflurane (3%) anesthesia.

### Aggressor Screening and Resident-Intruder (RI) Test

Mice were screened using protocols adapted from previous studies [6, 7]. Briefly, cage tops were removed and replaced with Plexiglas covers to monitor trials. Before initiating trials with paired female mice, the cohabiting male mouse was removed to a holding cage until completion of test. A novel C57BL/6J mouse matching the sex of the resident was introduced into each cage and mice were allowed to freely interact for 5-min. After 5-min elapsed, intruder mice were returned to their home cages and, in the case of female resident-intruder trials, cohabiting male mice were returned to their home cages. All videos were recorded for later analysis. Resident behaviors from the Mount Sinai videos were manually annotated using Observer XT 11.5 (Noldus Interactive Technologies). Behaviors annotated include anogenital (AG), chemoinvestigation, allogrooming, bite, facial chemoinvestigation, flank chemoinvestigation, kicking, lunging, pinning, withdrawal (active termination of interaction), “end” (passive termination of encounter) and wrestling.

### Behavioral definitions

#### 1. Investigation

Anogenital Chemoinvestigation: Resident mouse sniffs the anogenital region of the intruder.

Allogrooming: Resident grooms the back of the intruder.

Bite: Resident closes jaw on the body of the intruder.

Facial Chemoinvestigation: Resident mouse sniffs the face of the intruder.

Flank Chemoinvestigation: Resident mouse sniffs the flank of the intruder.

#### 2. Aggression

Kicking: Resident lifts hindleg to come in contact with the intruder.

Lunging: Resident propels itself at the intruder.

Pinning: Resident places paws on back of the intruder preventing the movement of the intruder.

Wrestling: Resident lunges toward the intruder and tumbles around the cage.

#### 3. Cessation of encounter

Withdrawal (active termination of interaction): Resident actively orients itself away from the intruder and walks away.

End (passive termination of encounter): Resident walks past the intruder without orienting away from the intruder or the intruder terminates the interaction.

### Aggression Conditioned Place Preference (CPP)

CPP testing was conducted in three phases as previously reported [7, 9]: pre-test, acquisition, and post-test. Mice were habituated to test rooms 1 hour before acquisition or test trials. All phases were conducted under red light and in sound-attenuated conditions. The CPP apparatus (Med Associates) has a neutral middle zone that allowed for unbiased entry and two conditioning chambers with different walls and floors. On the pre-test day, mice were introduced into the middle chamber and allowed to freely explore in all three chambers of the CPP box for 20 min. No group differences in bias for either chamber was found, and conditioning groups were balanced in an unbiased fashion to account for start side preference as described previously. The conditioning phase consisted of three consecutive days with two conditioning sessions each day, one in the morning (between 0900 and 1200 EST) and one in the afternoon (between 1400 and 1700 EST). During one session, experimental mice were confined to the assigned chamber for 10 min with a novel same-sex C57BL/6J intruder (paired session); during the other session, mice were put into the opposite chamber without a social target for 10 min (unpaired session). Aggression interaction was confirmed by an experimental observer to occur in the CPP cage. Timing of paired and unpaired sessions in the morning or afternoon was counterbalanced across the three-day acquisition period. On post-test day, experimental mice were placed into the middle chamber of the CPP apparatus and allowed to explore all chambers again freely for 20 min. Duration spent within either context was used to measure CPP. Behavioral analysis of CPP data was performed by assessing (*i*) subtracted CPP (post-test phase duration spent in the intruder-paired chamber minus test phase duration spent in the intruder-unpaired chamber, accounting for test session behavior only), and (*ii*) group and individual durations in both pre-test and post-test sessions.

### Aggression SA Apparatus

Aggression SA testing was conducted as described in .(Golden, et al., 2017, Golden, et al., 2019) Briefly, CFW resident mice were placed in standard Med Associates operant chambers and underwent 12 trials in which they could press a lever (FR1) in order to receive a subordinate same-sex mouse through a guillotine door into their operant chamber. A houselight illuminated the chambers during trials, and an inactive lever was extended at all times. At the beginning of a trial, the houselight turned on, and a lever extended 10s later. If the resident pressed for a reward, a 2s tone played and a guillotine door next to the active lever opened for 12s. A same-sex C57BL/6J intruder was ushered through the door from a custom 3d printed cannister, and the intruder was removed at the end of the trial. If residents did not press, the active lever retracted after 60s. All chambers also had recessed food pellet receptacles with beam-break registered entries ports.

### Food SA apparatus

Food SA testing was conducted in the aggression chambers using the opposite side houselight and lever, as well as a small cue light above the left-side active lever. Mice were tested for one hour and could receive up to 300 pellets per one-hour session on an FR1 schedule with a 20s timeout.

### Appetitive aggression SA

All mice underwent three consecutive days of resident intruder testing as described in. During RI training, mice were scored as either a 1 (non-aggressive), 2 (less than 5 seconds of aggression), or 3 (more than 5 seconds of aggression). Following RI training, each group was further separated into mice who were either AGG or NON during RI testing. Only one male was NON, and as such only AGG males were tested in operant SA. 3-d following RI testing, mice underwent 1-d of magazine training, in which they were exposed to operant cues (house light and a two second tone) in addition to a same sex intruder mouse entering the operant chamber (3 times each). On the following day, mice underwent SA training every other day for 9-d as previously described (Golden et al., 2017). Researchers were present throughout all aggression testing to ensure that no mice were injured. Mice with an average of 3 presses or less across days 4-8 of training were considered non-acquirers.

### Food SA

To control for learning ability in aggression non-acquiring mice, 9 male and 9 female aggression SA non-acquirers were tested for food training acquisition 7 days following the end of aggression SA. Mice underwent one day of autoshaping in which a house light of a different color was used for trials, in addition to a novel conditioned stimulus cue light and active lever location. Following autoshaping, mice underwent one hour per day for 7 days of food administration (FR1 schedule for 20mg food pellet with 20s timeout) for a possibility of 300 pellets per session.

### Data Analyses

Resident-intruder behavior quantification, conditioned-place preference, and aggression SA data were analyzed in Graphpad Prism (GraphPad Software) using parametric two-way analysis of variance (ANOVA), followed by multiple comparisons using the Tukey test. For comparisons of aggressive behaviors between male and female AGGs, Welch’s corrected t-test was used. Aggression SA statistical tests can be found in Supplemental Table 4 which includes chi-squared tests for proportions, repeated measures one-way ANOVAs for the female NON groups, two-way ANOVAs for male v female AGG comparisons, and mixed-effects with Geisser-Greenhouse correction tests for isolated female NON data due to missing data points.

### Hidden Markov Model

The hidden Markov model was performed using the R package hmm.discnp. Each mouse’s behavioral observations were generated as a unique sequence. The expectation maximization algorithm was iterated to find the state-dependent probability distributions (emission probabilities), state transition probabilities, and the initial state probabilities. We tested a 2, 3, 4, 5, and 6 state model and used the Bayesian information criterion to select which model best fit the data and found that a 4-state model fit best. The Viterbi algorithm was used to assign hidden states to each behavioral observation. In order to compare the percentage of observations in each state between groups, a two-way ANOVA was used followed by Tukey post-hoc comparisons.

### Random Forest Classifier

The random forest classifier was built using the tidymodels library suite in R (Kuhn & Wickham, 2020). The dataset consisted of all trials that were filmed on day three of RI. We then fit a random forest model as implemented by the ranger package for the purpose of classification. We constructed four models which used 20%, 40%, 60%, and 80% of the data for training. The number of variables to consider at each tree split was tuned using 10-fold cross-validation and the parameter that led to the best performance as indicated by the accuracy of the prediction and the area under the receiver-operator curve (AUC_ROC). We constructed a learning curve for each of the models by calculating the F1 score, which is the harmonic mean of the precision and recall of the model. Variable importance measures were determined by extracting the gini impurity metric from the model.

## Results

### Gross characterization of social behavior in male and female mice

During all three days of RI testing, both male and female AGGs engaged in more aggressive behavior than their NON counterparts (Phenotype F_(1, 51)_ = 36.20, p < 0.0001, Male AGG vs. Male NON, p =0.0002, Female AGG vs Female NON, p = 0.001), and there were no differences in duration or the latency to attack between male and female AGGs (p > 0.05) (Figures 1B & 1C, Supplementary Figures 1A-F). There were no differences between groups in investigative behavior on Day 3 (Sex: F_(1, 51)_ = 2.947, p = 0.0920. Phenotype: F_(1, 51)_ = 0.6242 Figure 1D), or on Days 1 and 2 Supplementary Figure 1C & 1G). On day 3, but not days 1 or 2, male NONs displayed a shorter latency to investigate intruders (Phenotype × Sex interaction F_(1, 51)_ = 5.003, p = 0.0297, Male NON vs Male AGG, p = 0.0139, Figure 1E).

**Figure 1.**
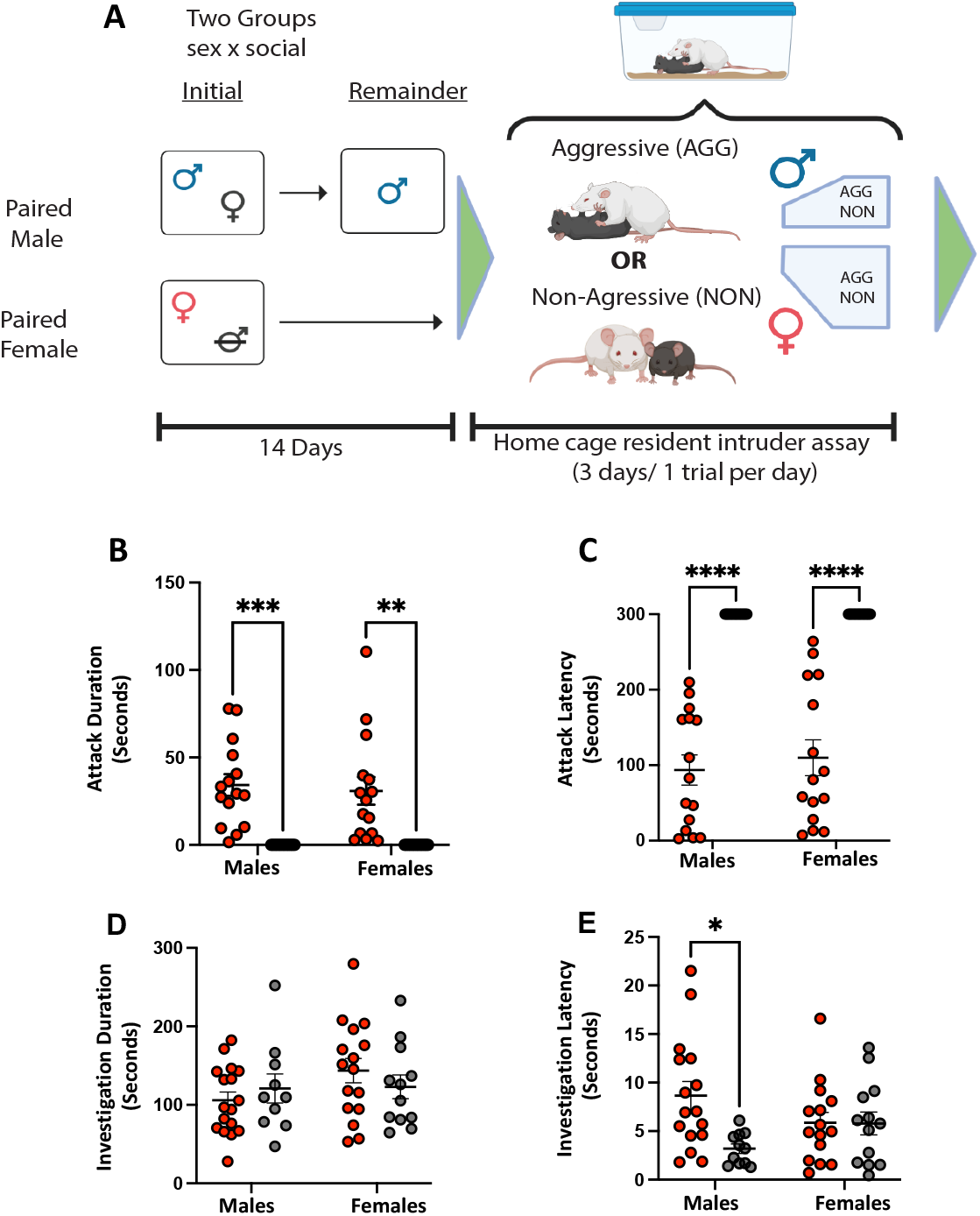
Male and Female SW mice engage in similar amounts of aggressive and investigative social behavior. (A). Schematic illustrating the housing conditions prior to the resident intruder test. (B). Total attack duration (B) and latency (C) did not significantly differ in male and female AGGs. (D). Total investigation. All groups show similar levels of social investigation E. Investigation latency. Male NONs had a significantly shorter latency to investigate the intruder than male AGGs.

### Male and female mice display distinct suites of social behavior

We next investigated whether male and female AGGs/NONs displayed distinct aggressive and/or investigative behaviors. We did not find any differences in anogenital (Sex: F_(1, 51)_ = 0.4110, p = 0.5243 Phenotype: F_(1, 51)_ = 1.141, p = 0.2905) or flank investigation (Sex: F_(1, 51)_ = 0.4182 p = 0.4551, Phenotype: F_(1, 51)_ = 0.0577, p = 0.8904) between any of the four groups on day 3 (Figure 2A & 2C) or on days 1 and 2 (Supplementary Figure 2A & Supplementary Figure 3A). Interestingly, we found that females engaged in significantly more facial investigation than males, regardless of their phenotype, on day 3 (F _(1, 51)_ = 10.54, p = 0.0021, Figure 2D). This effect was also seen on day 1 (Sex: F_(1, 52)_ = 8.751, p = 0.0046, Supplementary Figure 2D) with a trend towards significance (Sex: F_(1, 51)_ = 3.142, p = 0.0823) on day 2 (Supplementary Figure 3C). We also observed that AGGs, regardless of their sex engaged in more allogrooming than NONs (Phenotype: F_(1, 51)_ = 4.574, p = 0.0373), although only female AGGs were significantly different from female NONs (p = 0.0412, Figure 2B). Strikingly, on the initial day of the RI test, males did not engage in any allogrooming (Sex: F _(1, 52)_ = 34.01, p < 0.0001, Supplementary Figure 2B), while females did. Post-hoc comparisons once again revealed that female AGGs engaged in more allogrooming than their NON counterparts (p =0.0003). By day 2, males engaged in allogrooming at a level comparable to females. We also found a main effect of both sex (F_(1, 51)_ = 31.46, p < 0.0001) and phenotype (F_(1, 51)_ = 13.40, p = 0.006) on the number of withdrawals observed (Figure 3E), with female AGGs displaying a higher number of withdrawals than male AGGs (p < 0.0001) and Female NONs(p = 0.0022). No differences between the four groups were detected on days 1 (Supplementary Figure 2E) or 2 (Supplementary Figure 3E).

**Figure 2.**
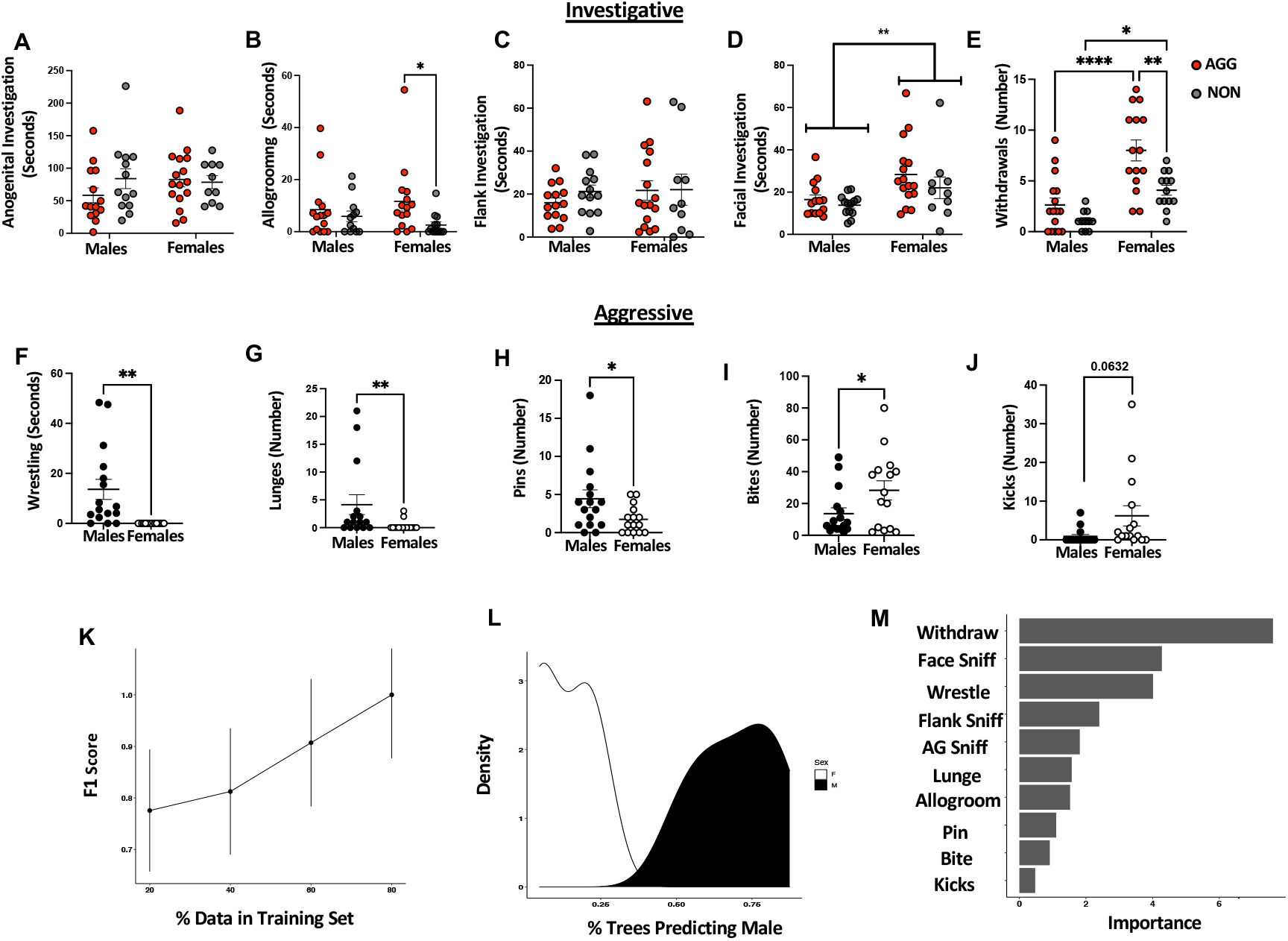
Male and female mice display distinct investigative and aggressive behaviors. For investigative behaviors, there were no group differences in anogenital investigation (A) or flank Investigation (C) AGGs regardless of sex spent more time allogrooming (B). Females regardless of phenotype engaged in more facial investigation (C) and withdrew from interactions more frequently (E). For aggressive behaviors, males engaged in more wrestling (F), lunges (G), and pinned (H) the intruder more than females. Females delivered more bites (I) and kicks (J) K. Learning curves from Random Forrest classifier. Curves were created using 1K trees, 4 data splits (20-80%), and with shuffled 10-fold cross-validation at each data split. Errors represent ± SEM. (L). Density plot demonstrating probability of being classified as M or F as a function of the number of trees predicting male. (M). Variable importance plot for the random forest classifier.

**Figure 3.**
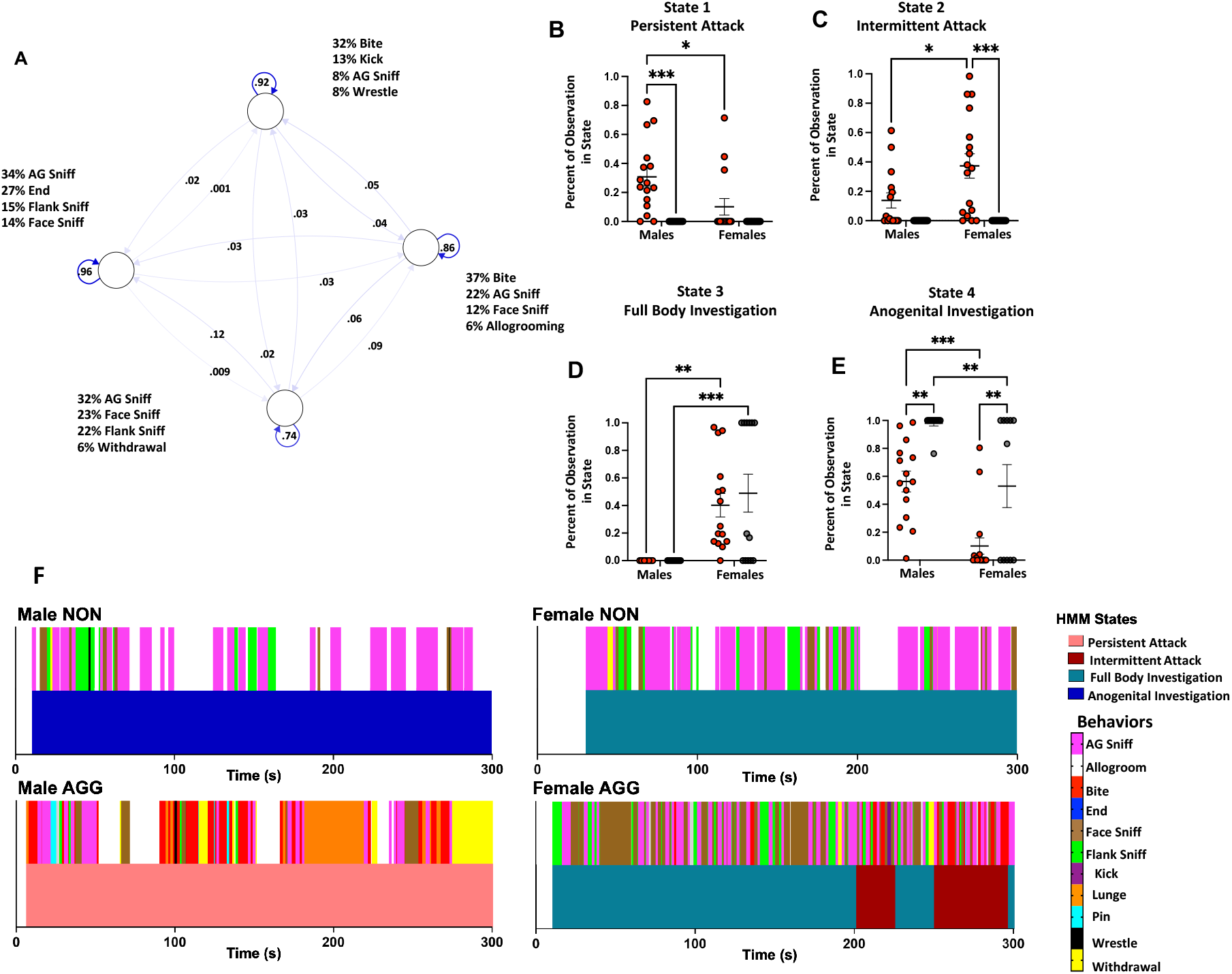
Hidden Markov Model of Social Behavior in the Resident Intruder Paradigm. (A). Schematic of HMM. Each node represents a hidden state. Numbers along the arrows indicate the probabilities of transitioning between states. Listed behaviors indicate the probability of occurrence during each state. Male AGGs were more likely to be in a state of persistent aggression (B) while female AGGs were more likely to be in a state of intermittent aggression (C). Females regardless of phenotype were more likely to be in the full-body investigation state than males(D). NON’s regardless of sex and males regardless of phenotype were more likely to be in the anogenital investigation state (E). (F) Representative examples of behavioral sequences (top) and predicted state (bottom) for all four groups.

When examining aggression, we observed that male and female mice engage in qualitatively distinct behaviors on day 3. Male AGGs engaged in wrestling behavior, in which the resident male lunges at the intruder and tumbles around the home cage, while female AGGs did not engage in this behavior at all (t(15) = 3.571, p = 0.0034, Figure 2F). Although some females did engage in lunging behavior, it was to a lesser extent than male AGGs (t(15) = 2.070, p = 0.054, Figure 2G). Females delivered more bites than males (t(15) = 2.104, p = 0.046, Figure 2I) with a and there was a trend for females to exhibit more kicks than males (t(15) = 2.005, p = 0.0632, Figure 2J). These single kicks were usually delivered following a single bite. In contrast, males were more likely to pin the intruder (t(15) = 2.151, p = 0.0442, Figure 2H) prior to delivering a bite.

Given that male and female mice display distinct sets of investigative and aggressive behavior, we used a random forest classifier to determine whether trials involving a male or a female as the resident were distinguishable based on the metrics quantified in Figure 2. Trials from day 3 were included in the model. We tested models in which 20, 40, 60, or 80 percent of the data was used for training the model (see Methods for details). We found that when 80% of the data was used to train the model, an F score of 1 was achieved, indicating a perfect classification of the remaining 20% of the trials (Figure 2K & L). We extracted the gini impurity metric to determine which variables were important for classifying males vs. females. The analysis indicated that withdrawals, facial investigation, and wrestling were important in classifying male vs. females.

### Male and female mice display distinct sequences of social behavior

For the HMM, we found that a 4-state model best fit the sequences of observations (see Methods for details). Inspection of the emission probabilities (Supplementary Table 1) suggests that states 1 and 2 (Persistent Attack & Intermittent Attack listed below as A1-A2 or I1-I2) were predominantly associated with aggressive actions, with bite being the most likely behavior to occur when the animal was in these states. Interestingly, state A2 was also characterized by a relatively high probability of investigation occurring, while state A1 was associated with relatively low probabilities of investigation (39% for state 2, 19% for state 1). Conversely, States 3 and 4 (Full Body Investigation & Anogenital Investigation) were predominantly associated with investigative behaviors, with aggressive behaviors being highly unlikely to occur (6% and .06% respectively). These investigative states were differentiated by the probability of specific investigative behaviors occurring. While in state I1, there was a roughly equal probability of anogenital (32%), facial (23%), and flank investigation (22%) (Supplementary Table 1). However, while in the anogenital investigation state, the mice were much more likely to engage in anogenital investigation (34%) rather than facial (14%) and or flank investigation (15%) (Supplementary Table 1). To determine whether certain groups were more likely to be in a particular state, we calculated the percentage of behavioral observations that occurred in each state for each mouse. We found that male AGGs had a significantly higher percentage of their observations in the persistent attack state than female AGGs (Sex x Phenotype interaction F_(1, 51)_ = 4.556, p = 0.0376, Male AGG vs. Female AGG, p =0.0111, Figure 3B).

Conversely, female AGGs had a significantly higher percentage of their observations in the intermittent attack state compared to male AGGs (Sex x Phenotype interaction F_(1, 51)_ = 4.451, p = 0.0398, Male AGG vs. Female AGG, p = 0.0206, Figure 3C). The difference between male and female AGGs is likely due to the fact that females are more likely to investigate the intruder before or after delivering a bite (36%) compared to males (14%) (Supplementary Table 3A & B) With regard to the full body investigation state, there was a striking sex difference, with none of the males showing any observations in this state (F_(1, 51)_ = 30.77 p < 0.0001, Male AGG vs. Female AGG, p = 0.0010. Female NON vs Male NON p =0.0023, Figure 3E). This phenomenon is due to the fact that females were more likely to string together multiple investigatory actions than males (Figure 3F, Supplementary Table 3A & B). Lastly, NON mice were more likely than AGGs to be in state I2, regardless of sex (phenotype F_(1, 51)_ = 25.85, p < 0.0001, Figure 3E).

### Socially housed males, but not females display appetitive aggression

In the CPP assay, there was a significant effect of time (F _(1,18)_ = 9.901, p = 0.0056). Post hoc analysis revealed that male AGGs but not male NONs spent more time in the paired chambered in the post-test relative to the pre-test (p < 0.05, Figure 4B). Neither female AGGs or NONs displayed a preference for the paired side during the post-test (Figure 4C).

**Figure 4.**
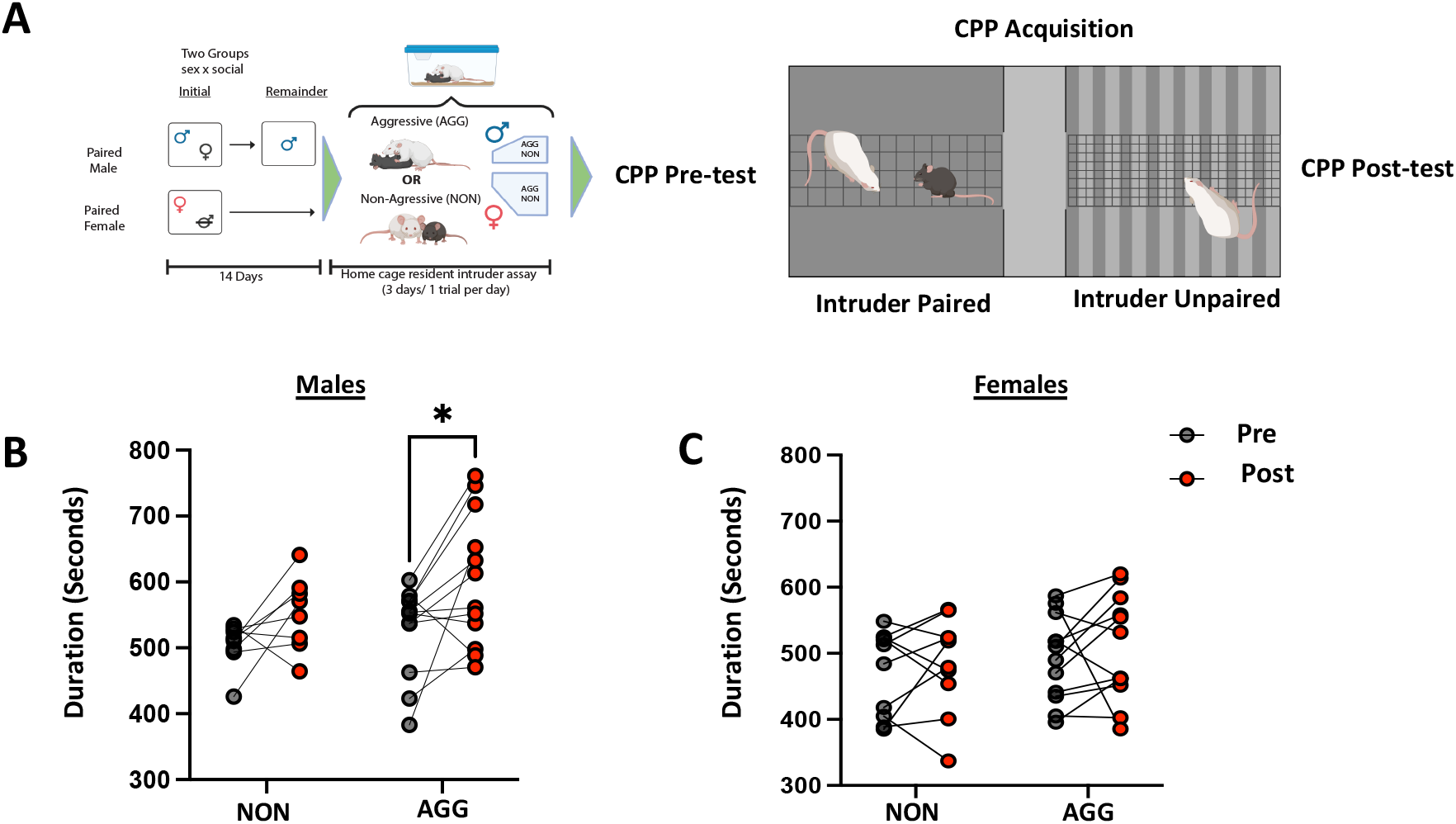
Male but not Female AGGs develop CPP to social encounters. A. Schematic of CPP paradigm. B) Male AGGs but not NONs develop a CPP to the paired chamber. C) Neither female AGGs or NONs developed a CPP to the paired chamber.

#### RI screening of paired aggression SA mice

Significantly more males then females were aggressive during RI screening (p < 0.001, Chisquare, df = 13.57,1, n = 29/29 AGG male, n = 18/29 female). The three resulting experimental groups (male AGG n= 9, female AGG n = 10, female NON n =7) differed significantly in latency to attack on day 3 of screening, with NON mice showing significantly longer latency to attack than the AGG groups (F _(2, 56)_ = 14.47, p < 0.0001, Figure 5B).

**Figure 5.**
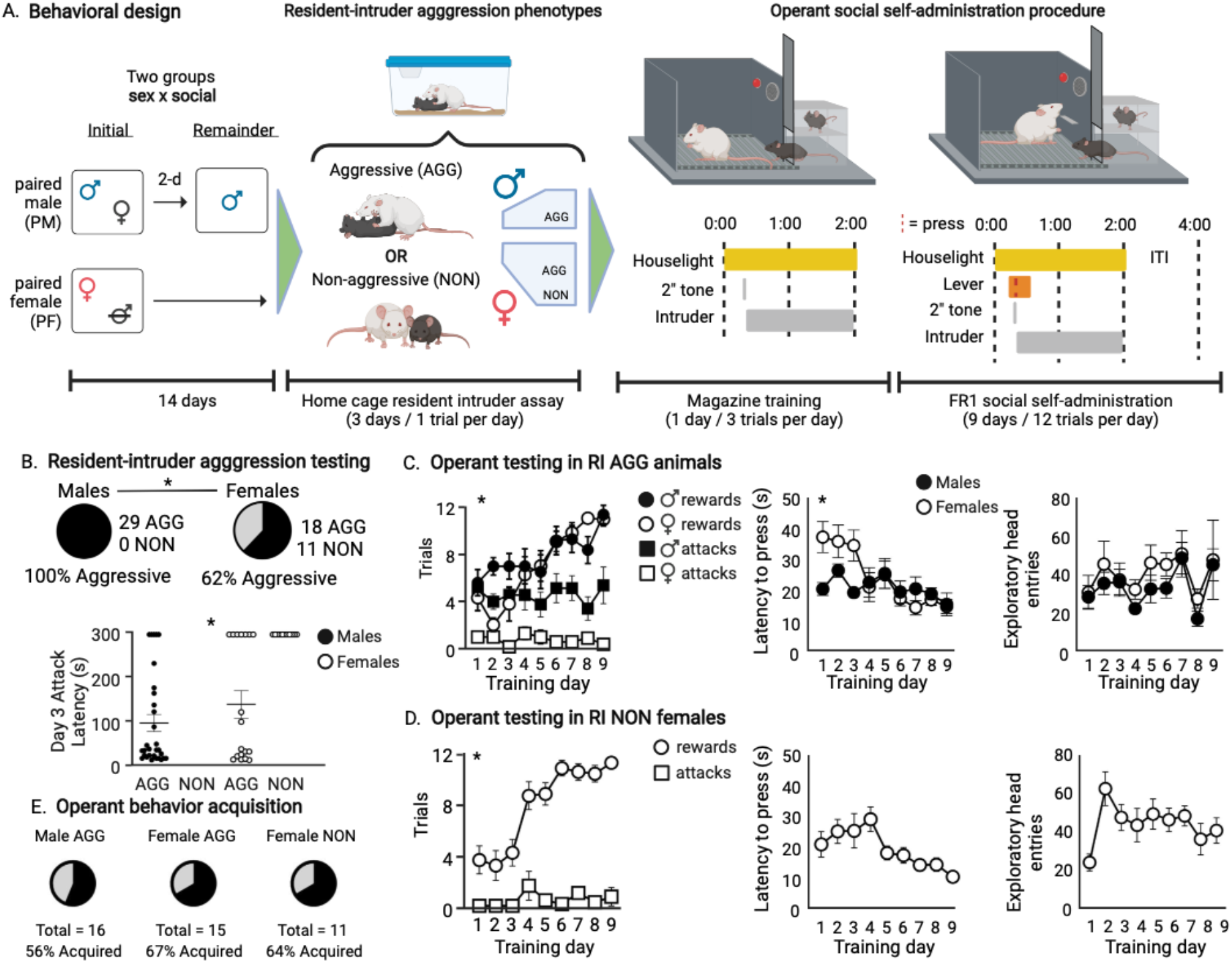
Males and females demonstrate similar aggression behavior in resident intruder but not appetitive aggression tasks. A) Schematic of social housing and behavioral paradigm. All males tested were aggressive during at least one trial of resident intruder screening, while the females separated into aggressive (AGG) and non-aggressive (NON) phenotypes. B) Latency to attack in the resident intruder assay differed significantly between groups, with female NONs having significantly higher latency to attack than the male or female AGGs. C) Females show slightly slower learning curves than males in acquiring the aggression self-administration task. Additionally, females show almost no attacks once they have self-administered a same-sex conspecific, while male aggression was steady across days. Females are initially slower than males to lever press, but both groups decrease latency over time. There were no differences in exploratory head entries across days or sex. D) Females who were not aggressive in the resident intruder screening show increasing rewards over time with steady attacks and decreasing latency to lever press. They show an increase in exploratory head entries initially which is steady thereafter. E) Similar percentages acquired operant selfadministration across groups.

#### Males and female AGGs learn to self-administer intruders similarly, but vary in attack behavior

There was a significant sex x day interaction in reward and attack behavior (interaction F_24,306_=3.327, p < 0.001, day F_8,306_ = 6.787, p < 0.001, sex F_3,306_ = 108.2, p < 0.001) with females showing significantly fewer attacks than males (p < 0.001, df = 306 Tukey’s).

Mice that did not acquire self-administration behavior were excluded from analysis. Latency to press for an intruder significantly decreased over days in both males and females (Interaction F _(8, 148)_ = 2.475 p = 0.015, Day F_8,148_=4.657 p < 0.001 Figure 5C), and there was no difference in exploratory head entry activity across days or sex. (F _(8, 153)_ = 0.1520, p = 0.9963, Figure 5C).

Female NONs showed similar lever press behavior compared to males, with significantly more rewards over time however, they exhibited significantly less (near zero) attacks across time (Interaction F_8,108_=9.277, p < 0.001, Day F_8,108_=13.02, p < 0.001, Attack v Reward F_1,108_=490.9, p < 0.001, Figure 5D). There were no significant differences across days in latency to press (F_(1.942,11.65)_ = 2.658, p = 0.11, Figure 5D) or exploratory head entries (F_(3.977,23.86)_ = 2.46, p = 0.07, Figure 5D).

There was no significant difference in the proportion of mice per group that acquired operant SA, as evidenced by an average of > 3 presses per day for the last five days of training (Male AGG = 9/16, Female AGG = 10/15, Female NON = 7/11, Chi-square, df = 0.3752, 2, p = 0.829 Figure 5E).

#### Housing condition does not impact appetitive aggression

Male and female AGGs that were housed in isolation for two weeks prior to the start of testing show similar trends to pair housed AGGs. In males (n = 8, Figure 6B), there was a trend toward increased rewards over time (F_2.487,17.41_=3.194, p = 0.057), with attacks remaining steady across days (F_2.557,17.9_=0.4350, p = 0.701). There were no significant differences in latency to press (F_3.892,26.27_ =0.833, p = 0.514) or exploratory head entries across days (F_2.224,15.57_=1.794, p = 0.197). In female AGGs (n = 9, Figure 6C) there was a significant increase in number of rewards across days (F_3.111,24.5_=7.995, p < 0.001), but no difference in number of attacks (F_2.571,22.82_=2.707, p = 0.0763). While latency to press decreased over time (F_2.746, 21.28_=5.199, p = 0.009), there was no change in number of exploratory head entries (F_3.134, 27.42_=1.444, p = 0.251).

**Figure 6.**
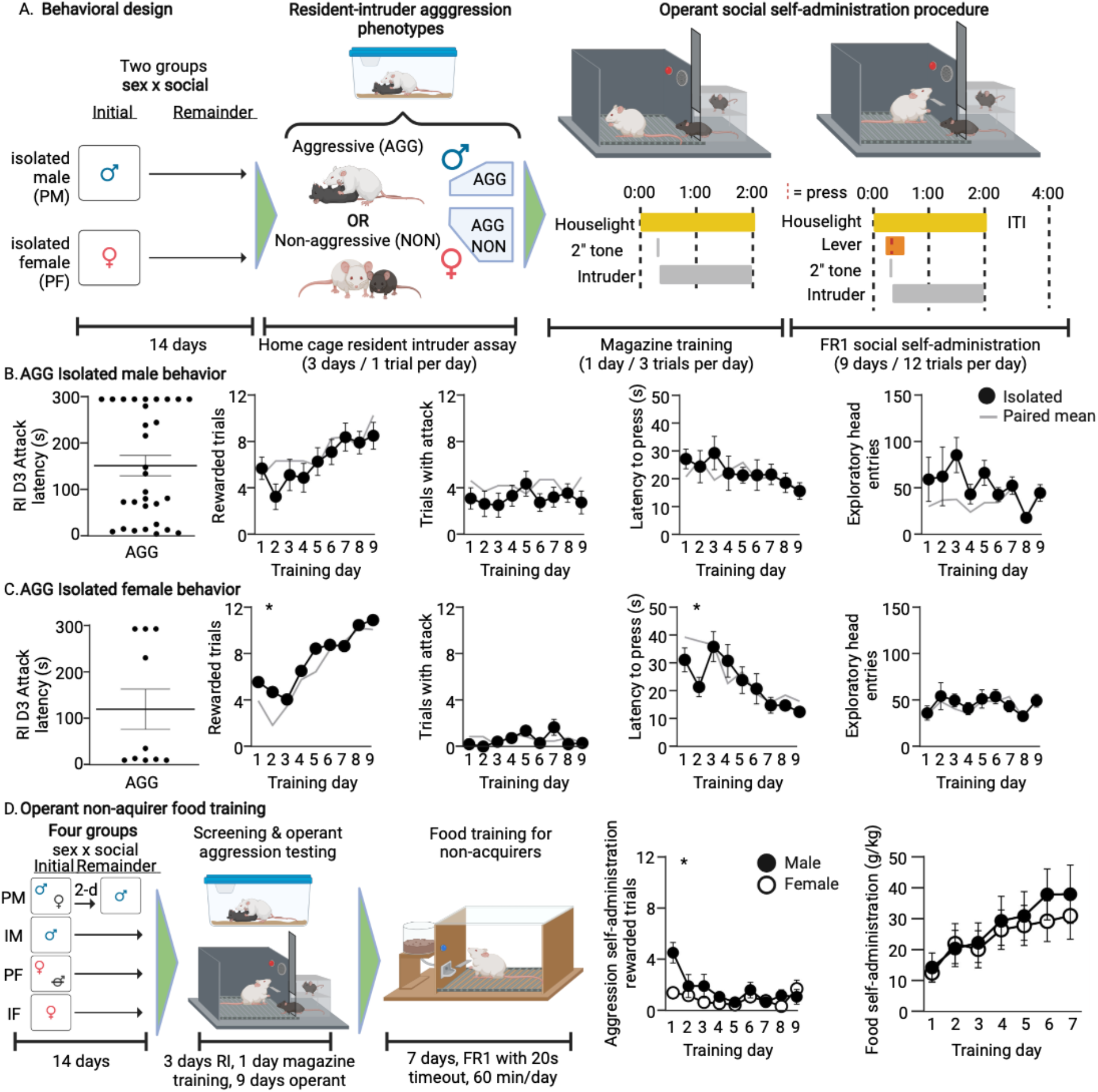
Isolate housing does not shift aggression patterns, and aggression self-administration non-acquirers rapidly learned food self-administration. A) Schematic of isolate housing and behavioral paradigm. B-C) Isolated males and females (black circles) show similar trends as socially housed mice (full data in figure XX, means showed here in gray). D) Abbreviated behavioral schematic showing housing conditions, resident intruder and aggression self-administration tasks, followed by seven days of sucrose pellet self-administration training for aggression non-acquirers. Males and females showed low rewards in aggression self-administration, with males initially slightly higher than females. Both males and females rapidly acquired sucrose pellet self-administration.

#### Aggression SA non-acquirers learn to self-administer palatable food

A subset of male and females (n = 9 male, 9 female) that did not acquire aggression self-administration were tested for learning capability via food self-administration testing (Figure 6D). There were no sex or housing differences in food self-administration performance (F _(6, 49)_ = 0.7440, p = 0.6169), and we therefore collapsed housing conditions across sexes for analysis. There was no sex x day interaction (F_6,96_=0.4544, p = 0.84) as both males and females similarly acquired the behavior.

## Discussion

We sought to characterize differences in aggressive and investigative social behavior in outbred male and female CFW mice. To this end, we adopted the protocol of Newman et al. [26] which allowed us to quantify aggressive social behavior in both sexes. Until now, most studies of female aggression in laboratory mice have resorted to using lactating females during the postpartum period [16, 17, 19, 27]. This is not ideal for evaluating sex differences in aggressive behavior since these behaviors are linked to hormonal changes specifically associated with pregnancy, parturition and lactation. Using this protocol, we found that when grossly measured as “aggressive” or “investigative”, males and females were largely similar although they exhibited sex specific behavioral sequences during bouts of aggression and investigation.

When rodents approach and contact a conspecific they engage in sniffing behavior of distinct body parts such as the face, anogenital, and flank regions [28]. We observed that females, regardless of their RI phenotype, engaged in facial investigation for longer durations than males. The facial area contains different excretory glands that give off distinct signals to the investigating animal. The Harderian glands are located near the eyes and excrete a lipid containing porphyrins [29] and have been shown to provide information about the sex and reproductive status of the individual [30]. The lacrimal glands are also located near the eyes and secrete fluid. These fluids have been shown to contain peptides such as extraorbital exocrine protein 1 (ESP1) which can influence social behavior in males [31, 32], but the effects certain facial cues have on female same-sex social interaction is unknown. Given that female AGGs spent significantly more time investigating the face of intruders relative to male AGGs, it is possible that specific chemical cues emitted from the face of females may promote aggression in female-female encounters, but this needs to be formally tested.

Males and females also displayed distinct attacking behaviors despite engaging in these behaviors for similar amounts of time and with similar latencies to onset. When male residents attacked the intruder, they displayed full-body lunges and wrestling behaviors that involved the two mice tumbling around the cage at very high speeds. This is in contrast to females, who were more likely to deliver a series of bites followed by a single kick with their hindlimbs. Male aggression thus seems much more explosive and offensive whereas female aggression seems tamer and possibly defensive in nature. This is in line with a previous study which investigated sex differences in attacking behavior in rats (Blanchard 1984). In that study, it was found that male bouts were more contact oriented with the male intruder having a higher chance of getting wounded from the bout relative to female intruders. In contrast, females were more likely to attack with a single bite or “jump-attack” (likely similar to the single kicks described in this paper) followed by the resident withdrawing from the encounter.

In addition to quantifying the duration of behaviors in male and female mice, we also employed a discrete state HMM. Although Markov chains have been used to examine aggressive behavior in males in the past [33, 34] this is the first instance of a hidden state model being used to analyze aggressive behavior in male and female mice to quantify sex differences. The discrete state HMM allowed us to go beyond a simple duration analysis by examining the temporal composition of social behaviors, and by clustering particular sequences of behavior into “hidden states”. We found that a 4-state model best fit our behavioral observations. Of these 4 states, states 1 and 2 were dominated by aggressive behaviors while states 3 and 4 were dominated by investigative behaviors. Interestingly, the “aggressive states” could be further differentiated by the probabilities of particular behaviors occurring. Although both states 1 and 2 were associated with a high level of aggression occurring, only state 2 was associated with a high level of investigation also occurring. This suggests that state 1 is characterized by persistent attacking for prolonged periods of time, while state 2 is characterized by a mix of both investigative and aggressive actions. Given the above discussion regarding the qualitative differences in attack behavior in males and females, it is not surprising that male AGGs had a greater proportion of their behaviors in state 1 whereas female AGGs had a greater proportion of their behaviors in state 2.

As with states 1 and 2, states 3 and 4 can also be further differentiated based on which particular behaviors were more likely to occur. State 3 was characterized by a roughly equal probability of any of the three main investigatory behaviors occurring, while state 4 was also characterized by a relatively high probability of AG investigation occurring relative to other modes of investigation. Strikingly, none of the behavioral sequences demonstrated by males were characterized as being in state 3. This is likely due to the fact that females were more likely to string together multiple investigative behaviors in succession, while males predominantly engaged in AG investigation or ended the interaction and then re-engaged in AG investigation during a separate bout. In contrast males tend to engage in interaction bouts that consist solely of one of the two types of social behavior (aggressive or investigative), terminate the bout, and then re-engage in a separate bout.

Although male and female AGGs displayed robust levels of reactive aggression, they differed with regard to aggression reward and the acquisition of appetitive aggression. The CPP experiment revealed that only male AGGs developed a preference for the side paired with aggressive experience, suggesting they find it to be rewarding or reinforcing. In line with these findings, while both males and females acquired SA behavior, only males attacked during the subsequent social interaction bout with the intruders. We can speculate that the robust female social self-administration may be affiliative, rather than aggressive, when social interactions are volitional rather than forced. These data agree with recently published work using outbred CD1 female mice, where female mice readily lever press for sensory contact to female partner mice [35]. However, our data also caution against the use of purely barrier-based social self-administration procedures in males and females due to the potential incongruence in aggressive behavior between RI and SA testing. Use of barrier and purely sensory contact may mask, whether aggressive or affiliative, the ultimate motivation of the resident mouse.

Interestingly, male CFW mice did not show an increase in attack trials across training days, which is a characteristic of appetitive aggression in male CD1 mice. As with inbred mouse strains, these data highlight the importance of the selection of either inbred or outbred lines for social behavior studies. Further, these results indicate that female CFW mice are a valid model for studying reactive aggression, which is a departure from the historical narrative that female mice are only maternally aggressive and can therefore be excluded under the NIH sex as a biological variable initiative. Female CFW mice cannot, however, be used to examine appetitive aggression behavior using operant self-administration procedures.

Several other considerations were explored in this study and warrant further discussion. First, we found no impact of housing condition on aggression in the RI or operant SA test, indicating that female CFW mice do not necessarily need to be primed by cohousing with castrated males to demonstrate reactive aggression, and can instead undergo isolate housing as is the typical procedure for males. While we did see differences in the percentage of males that were NON versus AGG in RI testing between Mount Sinai and the University of Washington, this is not unexpected. Outbred lines, while helpful in studying individual differences in aggression, can exhibit batch differences due to the nature of their genetic variability, as has been seen in CD1 outbred mouse aggression testing (Kwiatkowski et al., 2021). Additional differences could be explainable due to differences in protocols included the vendor in which CFW mice were purchased, screening during the light/dark cycle, age of intruder, and sex of experimenters, any or none of which may have been responsible for these differences. As such, variation between sites in percentage of aggressive CFW mice during RI testing is not unexpected and we recommend each site establish baseline aggression profiles in both males and females using the RI test before implementing their experimental protocols.

In summary, we show that despite similar levels of aggression and investigation, the actions displayed by male and female residents—which make up the gross measures of social behavior—are both qualitatively and quantitatively distinct. Our HMM revealed that females are more likely to switch between aggressive and investigative behaviors within a given interaction bout, while males typically engage in only one of these behaviors per bout. Furthermore, while female outbred CFW mice exhibit reactive aggression, only male outbred CFW mice displayed robust levels of appetitive aggression in CPP and SA experiments. Thus, future studies to disentangle the underlying biology driving these sex differences are critical.

## Funding and Disclosure

The authors declare that they do not have any conflicts of interest (financial or otherwise) related to the text of the paper. The research was supported R01MH127820 (SJR), R01MH114882 (SJR), R01MH104559 (SJR), R01 MH120514 (SJR), R01 MH120637 (SJR), R00DA045662 (SAG), P30DA048736 (SAG), NARSAD Young Investigator Award 27082 (SAG), and F31MH125587-01 (NLG). Some figures created with BioRender.com.

---

**Figure S1.**
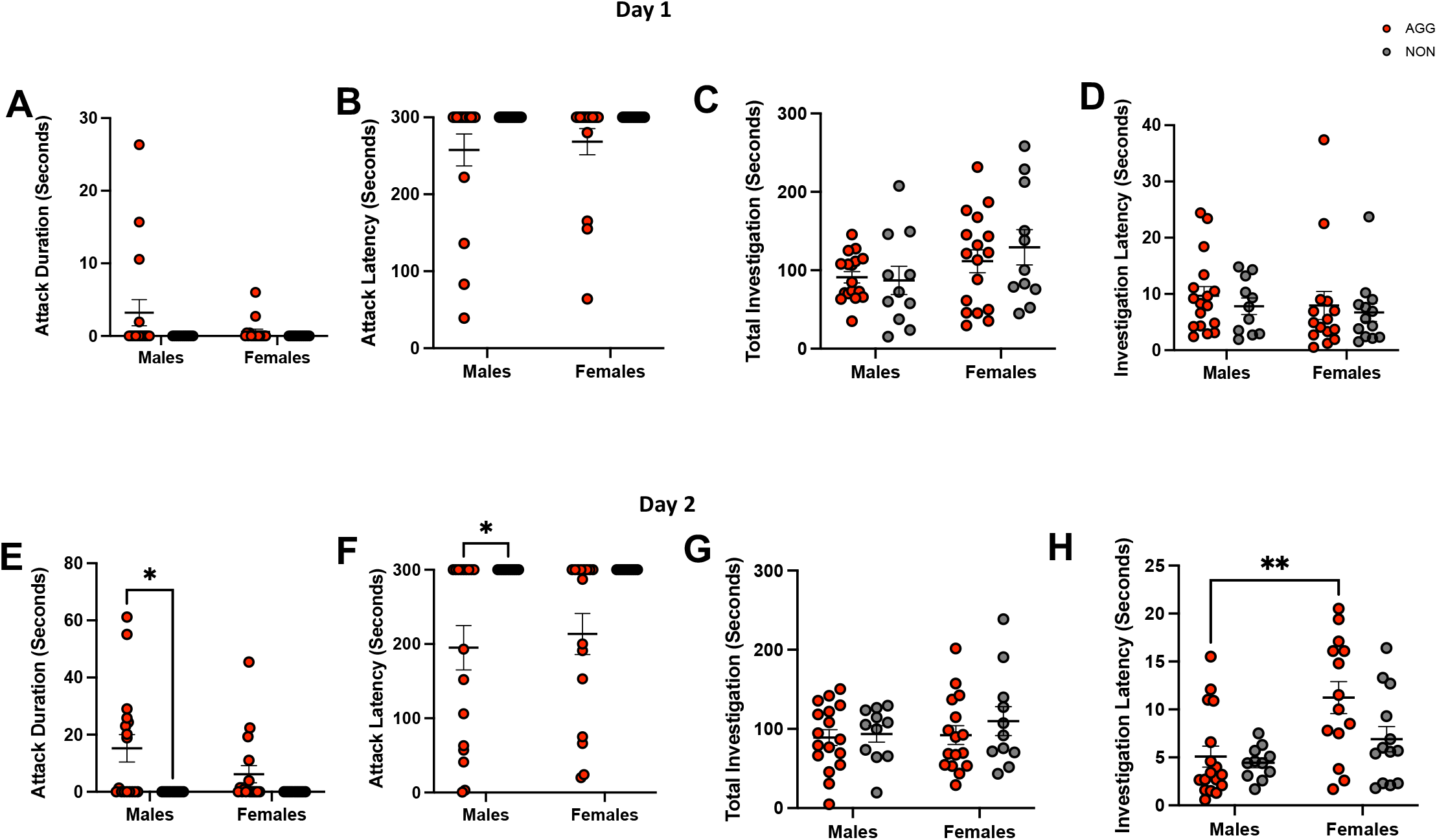
Gross measures of social behavior on Days 1 and 2. A. Attack duration. Twoway ANOVA interaction F (1, 52) = 1.320, p = 0.258. B. Attack Latency. Two–way ANOVA, main effect of phenotype F (1, 52) = 4.832, p = 0.0352. C. Total investigation. Two-Way ANOVA, main effect of sex, F (1, 52) = 4.178, p = 0.046. D. Investigation latency. Two-way ANOVA interaction F (1, 52) = 0.02575, p = 0.8731. E. Attack duration (Day 2). Two-way ANOVA, main effect of phenotype, F (1, 52) = 9.167, p = 0.0038. Tukey’s post-hoc Male AGG vs. Male NON, p = 0.0183. F. Attack latency (Day 2). Two-way ANOVA, main effect of phenotype F (1, 51) = 14.42, p = 0.004. G. Total investigation (Day 2). Two-way ANOVA, F (1, 51) = 0.2639, p = 0.6097. H. Investigation latency (Day 2). Two-way ANOVA, main effect of sex F (1, 51) = 11.34, p = 0.0015. Tukey’s post-hoc female AGG vs male AGG, p = 0.0035.

**Figure S2.**
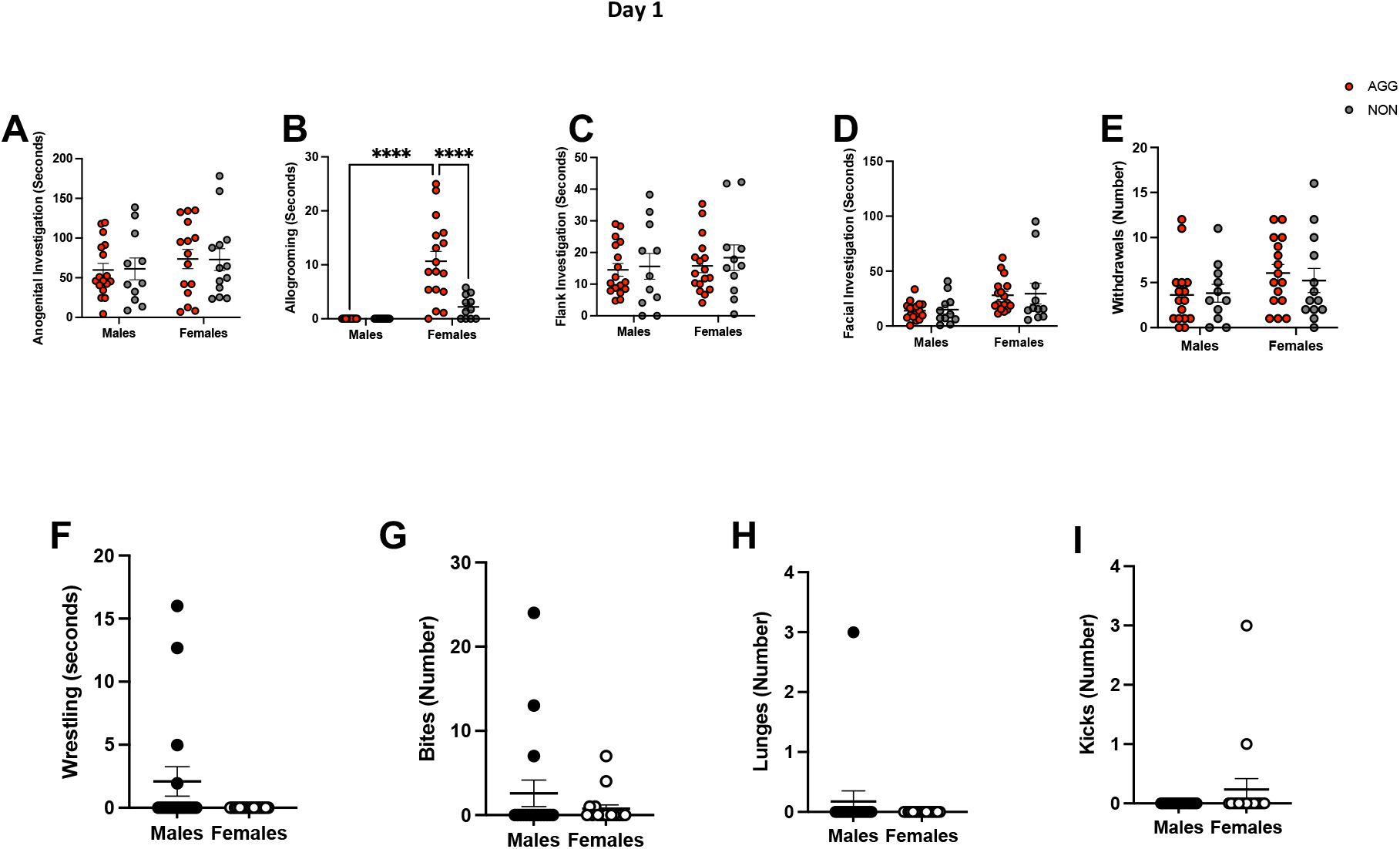
Quantification of distinct social behaviors on day 1. A. Anogenital investigation.Two-way ANOVA, no effect of sex or phenotype. B. Allogrooming. Two-way ANOVA, sex x phenotype interaction, F (1, 52) = 9.518, p = 0.0033. Tukey’s post-hoc test, female AGG vs male AGG, p < 0.0001. Female AGG vs female NON, p = 0.003. C. Flank investigation. Twoway ANOVA, no effect of sex or phenotype. D. Facial investigation. Two-way ANOVA, main effect of sex F (1, 52) = 8.751, p = 0.0046. E. Withdrawals, no effect of sex or phenotype F. Wrestling, Welch’s t-test t (16) = 1.793, p = 0.09. G. Bites, Welch’s t-test t (18.67) = 1.108, p = 0.281. H. Lunges, Welch’s t-test t (16) = 1.00, p = 0.3322. I. Kicks, Welch’s t-test t (16) = 1.289, p = 0.215.

**Figure S3.**
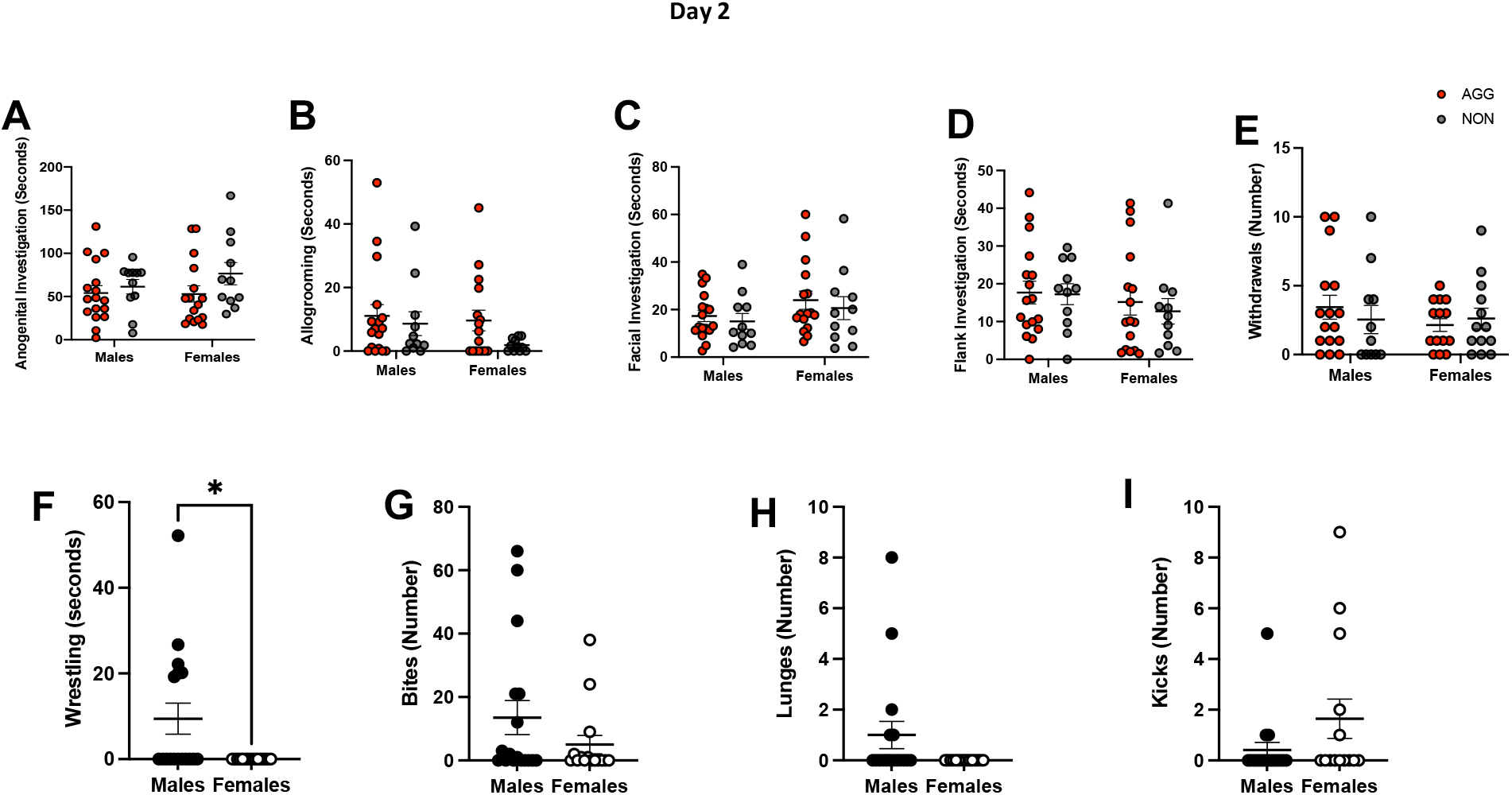
Quantification of distinct social behaviors on day 2. A. Anogenital investigation Two-Way ANOVA, no effect of sex or phenotype B. Allogrooming. Two-way ANOVA, no effect of sex or phenotype. C. Flank investigation. Two-way ANOVA, no effect of sex or phenotype. D. Facial investigation. Two-way ANOVA, main effect of sex F (1, 51) = 3.142, p = 0.0823. E. Withdrawals. Two-Way ANOVA no effect of sex or phenotype. F. Wrestling, Welch’s t-test t (16) = 2.59, p = 0.0194. G. Bites, Welch’s t-test t (24.22) = 1.402, p = 0.1735. H. Lunges, Welch’s t-test t(16) = 1.867, p = 0.0803. I. Kicks, Welch’s t-test t (16.83) = 1.483, p = 0.1565

**Supplementary Table 1.** Emission probability matrix. Each cell indicates the probability of observing a given behavior in each of the four states.

|  | S1 | S2 | S3 | S4 |
| --- | --- | --- | --- | --- |
| AG sniff | 0.08605625 | 0.22792224 | 0.32384628 | 0.34003168 |
| Allogrooming | 1.39E-26 | 0.06597771 | 0.05617866 | 0.03629472 |
| Bite | 0.32261289 | 0.3721225 | 0.01189169 | 5.79E-06 |
| End | 0.11603446 | 0.02531174 | 0.06355925 | 0.27885629 |
| Face sniff | 0.08426891 | 0.12043818 | 0.2316754 | 0.14977904 |
| Flank sniff | 0.02677525 | 0.04650898 | 0.23597293 | 0.15385922 |
| Kick | 0.13225733 | 0.03338049 | 0.00245818 | 5.73E-62 |
| Lunge | 0.08537152 | 6.75E-18 | 1.31E-68 | 6.39E-14 |
| Pin | 0.02424978 | 0.05413276 | 0.00619664 | 0.00841112 |
| Withdraw | 0.04750985 | 0.05280663 | 0.06822098 | 0.03190471 |
| Wrestle | 0.07486376 | 0.00139876 | 9.10E-57 | 0.00085743 |

**Supplementary Table 2.** Transition probability matrix. Each cell indicates the probability of transition from one state to another.

|  | S1 | S2 | S3 | S4 |
| --- | --- | --- | --- | --- |
| S1 | 0.92266881 | 0.04333721 | 0.02364991 | 0.01034407 |
| S2 | 0.05223531 | 0.86446045 | 0.05780613 | 0.02549812 |
| S3 | 0.03172461 | 0.09440697 | 0.74857678 | 0.12529164 |
| S4 | 0.00236107 | 0.00951385 | 0.02598266 | 0.96214243 |

**Supplementary Tables 3A and 3B.** Transition probabilities between investigate behaviors (A) and between bites and investigative behaviors in males and females (B). Key: AG: anogenital, B: Bite, FA: Face, FK: flank.

|  | M | F |
| --- | --- | --- |
| FA->AG | 0.32 | 0.46 |
| FA->FK | 0.2 | 0.14 |
| FK>FA | 0.08 | 0.25 |
| FK>AG | 0.34 | 0.51 |
| AG>FA | 0.06 | 0.18 |
| AG>FK | 0.19 | 0.3 |
| B->AG | 0.09 | 0.26 |
| B->FA | 0.01 | 0.10 |
| B->FK | 0.04 | 0.04 |
| AG->B | 0.05 | 0.21 |
| FA->B | 0.05 | 0.08 |
| FK->B | 0.04 | 0.03 |

**Supplementary Table 4.**
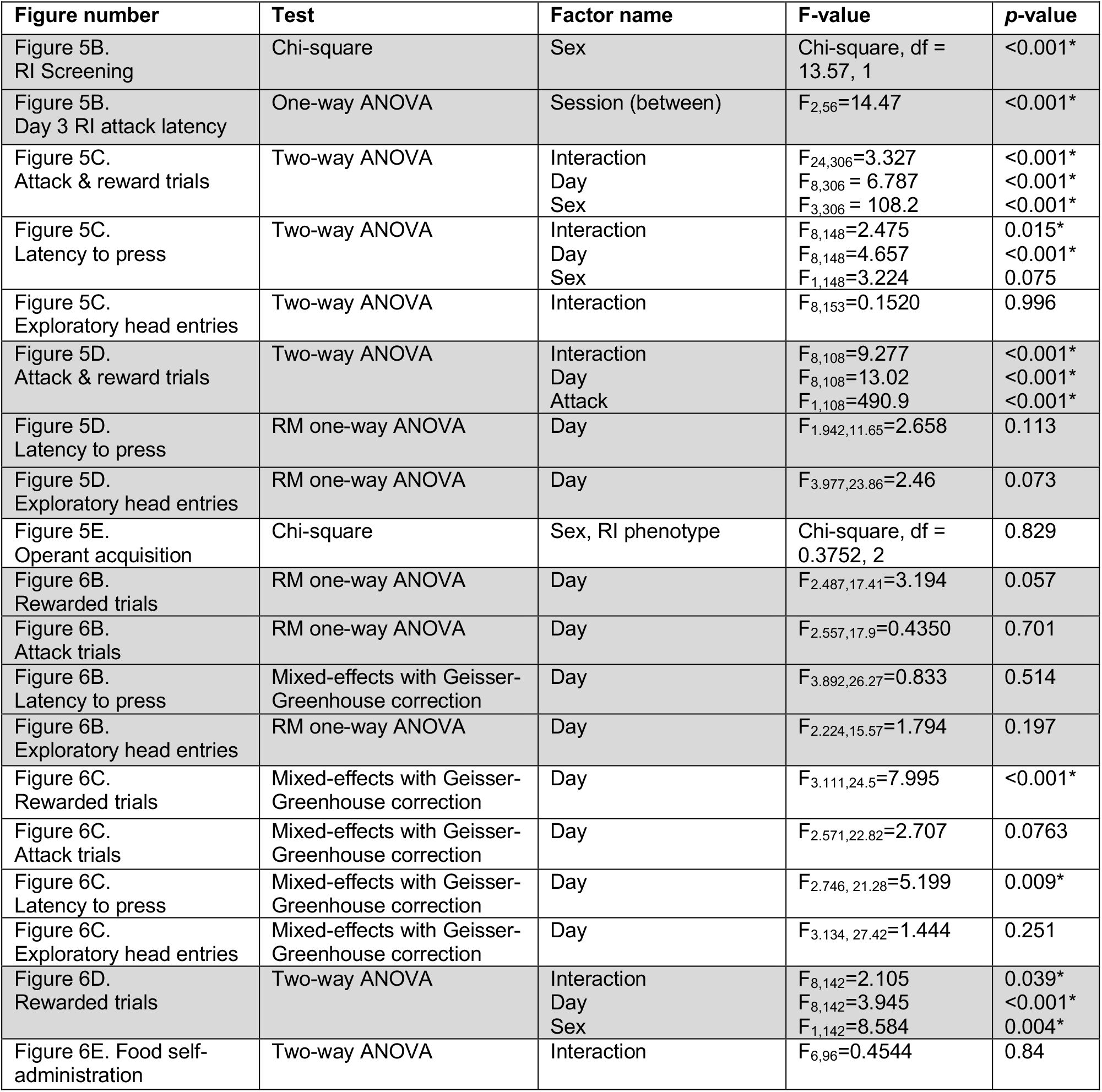
Data analysis from self-administration experiments

## Notes

### Competing Interest Statement

The authors have declared no competing interest.

